# Low exposure to lithium does not induce nephrogenic diabetes insipidus but microcystic dilations of collecting ducts in a long-term rat model

**DOI:** 10.1101/2023.10.12.562078

**Authors:** Nahid Tabibzadeh, Mathieu Klein, Mélanie Try, Joël Poupon, Pascal Houillier, Christophe Klein, Lydie Cheval, Gilles Crambert, Samia Lasaad, Lucie Chevillard, Bruno Megarbane

**Author notes:** **Correspondence:** Laboratoire de Physiologie Rénale et Tubulopathies, Centre de Recherche des Cordeliers, Inserm, Sorbonne Université, Université Paris Cité, Paris, France. *E-mail address:* (Dr. N. Tabibzadeha) and Réanimation Médicale et Toxicologique, Hôpital Lariboisière, Fédération de Toxicologie, APHP, Paris, France. *E-mail address:* (Dr. B. Mégarbane). Authors contributed equally to the present work (Nahid Tabibzadeh, Mathieu Klein, Mélanie Try).

## Abstract

Lithium induces nephrogenic diabetes insipidus (NDI) and microcystic chronic kidney disease (CKD). As clinical studies suggested that NDI is dose-dependent and CKD time-dependent, we investigated the effects of low exposure to lithium in a long-term rat model. Sprague-Dawley rats were randomly fed during six months with normal diet (controls), addition of lithium to diet, or addition of lithium and amiloride to diet, allowing reaching low steady-state plasma lithium concentrations (0.25±0.06 and 0.43±0.16 mmol/L, respectively). Exposure to low plasma lithium concentrations did not induce NDI but microcystic dilations of kidney tubules, identified as collecting ducts (CDs) using immunofluorescent staining. Both hypertrophy, characterized by an increase in the ratio of nuclei per tubular area, and microcystic dilations were observed. Principal cell-to-intercalated cell ratio was higher in dilated than in hypertrophied tubules. There was no correlation between aquaporin-2 mRNA levels and cellular remodelling of CDs. Amiloride/lithium co-administration did not allow significant consistent morphometric and cellular composition changes compared to lithium administration. To conclude, rat low exposure to lithium did not induce overt NDI but microcystic dilations of CDs, which include a marked alteration in cell composition of hypertrophied and dilated CDs, suggesting two distinct underlying pathophysiological mechanisms.

## Introduction

Lithium is the first-line treatment of bipolar disorder patients (Geddes et al. 2004, 2010; Miura et al. 2014); however, its high efficacy is counterbalanced by long-term adverse effects (Tabibzadeh et al. 2019). Renal adverse effects are the most remarkable and could be divided in two categories, nephrogenic diabetes insipidus (NDI) and microcystic chronic kidney disease (CKD) (McKinght et al. 2012; Shine et al. 2015).

Regarding lithium-induced microcystic CKD, experimental studies have reported tubular dilations and microcysts, developing as soon as 1 month after lithium treatment initiation (Hestbech et al. 1977; Kling et al. 1984; Walker et al. 2013). Interstitial fibrosis and uremia have also been induced in long-term lithium exposure models up to 6 months (Christiensen et al. 1983; Walker et al. 2013). Based on morphological analyses and lectin staining, distal tubules and/or collecting ducts (CDs) have been indirectly identified as the involved nephron structures by dilations and microcysts (Walker et al. 2013; Markowitz et al. 2000). However, no investigation based on specific staining has been reported, and the exact cell composition of those cysts has not been clearly established. Experimental data showed that CD remodelling is characterized by cell proliferative signals and an increase in type-A intercalated cell (AIC) to principal cell (PC) ratio (Christiensen et al. 1983; Nielsen et al. 2008; de Groot et al. 2014).

Lithium-induced NDI is defined by early polyuria and polydipsia due to renal resistance to vasopressin secondary to a decrease in apical membrane addressing and expression of aquaporin-2 (AQP2) (Alsady et al. 2016). NDI is usually observed at lithium concentrations in the therapeutic range, starting at 0.5-0.6 mmol/L (Nielsen et al. 2006; Tabibzadeh et al. 2019). Whether a mechanistic relationship exists between lithium-induced NDI and CKD is unclear. Moreover, the relationship between microcystic tubular dilations, cell composition of microcysts, and resistance to vasopressin remains to be established.

NDI develops shortly after lithium treatment initiation in humans as well as in experimental models, while CKD is inconstant and slowly progressive (Walker et al. 2005; Davis et al. 2018). Recent clinical studies suggested distinct determinants of lithium-induced NDI and CKD. While the daily lithium dose is considered as the main NDI determinant (Tabibzadeh et al. 2022), lithium treatment duration is reported to be independently associated with CKD onset (Tabibzadeh et al. 2021).

Based on case series, amiloride has been proposed as a therapeutic option of lithium-induced NDI patients (Battle et al. 1985; Bedford et al. 2008). Amiloride is a specific blocker of the Epithelial Sodium Channel (ENaC), hence of lithium entry in PCs (Kortenoeven et al. 2009; Tabibzadeh and Crambert 2022). Regarding lithium-induced microcystic CKD, two studies have shown a reduction in kidney fibrosis and tubular dilations in rats receiving a 6- month daily lithium regimen at pharmacological doses if lithium was co-administered with amiloride (Kalita-De Croft et al. 2018; Mehta et al. 2022). Given these findings, we designed a rat study aiming i)- to investigate the effects of an under-exposure to lithium on NDI and microcystic tubular dilations; ii)- to clarify possible relationships between these two types of renal injury; and iii)- to evaluate the effectiveness of amiloride to prevent them.

## Materials and methods

### Animals

All procedures were performed in accordance with the French animal care legislation and approved by the Ethics committee of Paris Cité University and the French Ministry of Research (approval # 26674-2020020714451509). Wild-type male Sprague-Dawley rats (mean weight of 100-125 g, Janvier Labs) were fed using normal rat diet with free access to water. Rats were randomly allocated to three groups (n=8 /group): (1) fed with diet alone (control group), (2) fed with diet added with lithium (Li^+^ group), and (3) fed with diet added with lithium and amiloride (Li^+^/Ami group) for a 6-month period. Lithium carbonate (Sigma, Saint-Quentin-Fallavier, France) was added to the dry food at 0.1% (0.05% of wet food, corresponding to 2.7 mmol/kg of lithium). Amiloride hydrochloride (Gerda, Paris, France) was added to the rat diet at 2.5 mg/kg dose in the Li^+^/Ami group.

### Sample processing

Urine and plasma were collected at 4, 5, and 6 months during metabolic cage housing. Creatinine concentration measurement was performed using a Konelab™ device (ThermoFischer Scientific, Waltham, MA, USA). Urine osmolality was measured using the Advanced 3250™ osmometer (Advances Instruments, Norwood, MA, USA). Plasma Lithium was measured using Inductively Coupled Plasma Mass Spectrometry assay (NexION 2000™ spectrometer, Perkin Elmer, Villebon-sur-Yvette, France). At the end of the protocol (6 months), the left kidney was rapidly excised under anaesthesia, partly embedded in OCT and frozen for fluorescent immunostaining, and partly frozen and conserved at -80°C for subsequent RNA analyses. Subsequently, aorta was cannulated to infuse isotonic saline followed by 4% paraformaldehyde (PFA) in the right kidney, which was removed, fixed in 4% PFA solution, embedded in paraffin, and further processed for Masson trichrome staining (3µm sections).

### Urine concentration test

Water deprivation and desmopressin challenging tests were performed separately twice, after 3 and 6 months of treatment in metabolic cages. Water deprivation test consisted in a 6- hour hydric restriction with urine sampling at 2, 4 and 6 hours. Desmopressin challenging test consisted in intramuscular desmopressin (DDAVP, 0.75 µM/kg) administration after 2 hours of hydric restriction, with subsequent urine collection at 2 and 4 hours post-injection.

### Immunofluorescence assay

Snap-frozen kidney samples were processed after cryosection (4µm sections) for immunofluorescence microscopy using a two-step staining. In the first step, primary antibodies consisted in goat anti-AQP2 SC-9882, 1/500 (Santa Cruz, Biotechnology Inc., Dallas, Tx, USA) and rabbit anti-AE1 20112, 1/800 (Cell Signalling Technology, Danvers, MA, USA). Secondary antibodies consisted in donkey anti-goat AF488, SC-362255, (Santa Cruz, Biotechnology Inc., Dallas, Tx, USA) and donkey anti-rabbit AF555 A-31572, 1/500 (ThermoFischer Scientific, Waltham, MA, USA). In the second step, direct FITC-labelled Ki67 1/500 antibody were used (ab281847; Abcam PLC, Paris, France).

### Morphometric analyses

All sagittal sections of kidneys were scanned using Axioscan Z1 slide scanner (Zeiss, Oberkochen, Germany) after Masson trichrome staining or immunofluorescent labeling. Tubular area was semi-automatically measured using the QuPath software (Bankhead et al. 2017). Briefly, approximately one sixth of the cortex was selected on Masson-stained tissue slices, and vessels (arteries and veins) were manually excluded. A pixel classification method was trained from manual annotations, to differentiate tubular lumen and parenchyma. A cut-off of 1500 μm^2^ was then applied on ’lumen’ objects to exclude glomerulus urinary spaces and peritubular capillaries. An average of 436 ± 223 tubules were analysed in each kidney.

The ratio between PCs (stained with AQP2) and AICs (stained with AE1, anion-exchanger-1), and between number of cells and tubular area was assessed in dilated and non-dilated CDs.

### RT-PCR

Kidney samples were separated immediately after excision to obtain specific samples from the cortex, the outer stripe of the outer medulla, the inner stripe of the outer medulla, and the inner medulla (including the papilla). Total RNA was extracted from these samples using the RNeasy Micro-Kit according to the manufacturer’s protocol (Qiagen, Hilden, Germany). Reverse transcription was performed using first-strand cDNA synthesis kit for RT-PCR (Roche Diagnostics, Basel, Switzerland). qPCR was then performed on a LightCycler™ (Roche Diagnostics) using a SYBR™ green kit (LightCycler 480 SYBR Green Master, Roche Diagnostics) for AQP2 (forward primer: 5’-ACC TGG CTG TCA ATG CTC TC - 3’ and reverse primer: 3’-GG ACG GGA GAG GTA ACC AAA - 5’), normalizing to the ribosomal protein S23 (RPS23) transcript levels (forward primer: …, reverse primer:…).

### Statistical analyses

Data are presented as percentages or mean ± SEM values, as appropriate. Non-parametric Mann-Whitney or Kruskal-Wallis tests were performed to test differences between groups as requested. Non-parametric Spearman tests were used to test correlations between AQP2 expression and cellular quantifications. The significance level of a statistical hypothesis test was set at 0.05. All statistical analyses and graphs were performed using Prism GraphPad™ (Boston, MA, USA).

## Results

### Under-exposure to lithium resulted in no overt NDI but in microcystic dilations of CDs

Plasma lithium concentration in the Li^+^ group was 0.25 ± 0.06 mmol/L. Rats displayed non-significantly higher urine output but lower urine osmolality than controls (Fig. 1). Urine concentration challenging tests (hydric restriction and desmopressin injection) did not reveal significantly altered urine concentration ability in the Li^+^ group rats at 3 and 6 months of treatment. After 6 months of treatment, serum creatinine concentration did not significantly differ between the two groups, and no overt interstitial fibrosis was observed on histopathological assessment (Fig. 2). However, consistent and marked tubular morphological changes were observed in the Li^+^ group rats, characterized by cortical and outer medullary microcystic tubular dilations, sometimes associated with intratubular casts. Semi-automated tubular area measurement showed significantly higher tubular areas and minimal diameters in the Li^+^ group rats compared to controls (Fig. 2).

**Fig. 1.**
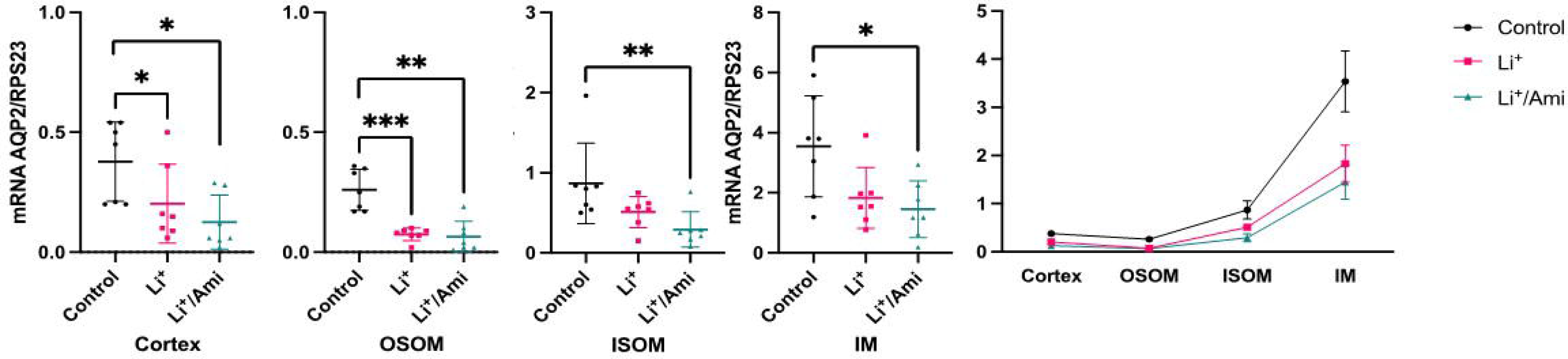
Urine output, urine osmolality, and serum creatinine concentration. Data are expressed as mean ± SEM. Uosm, urine osmolality; Li^+^, lithium; Li^+^/Ami, lithium and amiloride treatment; DDAVP, desmopressin. There were no significant differences between the groups in all analyses.

**Fig. 2.**
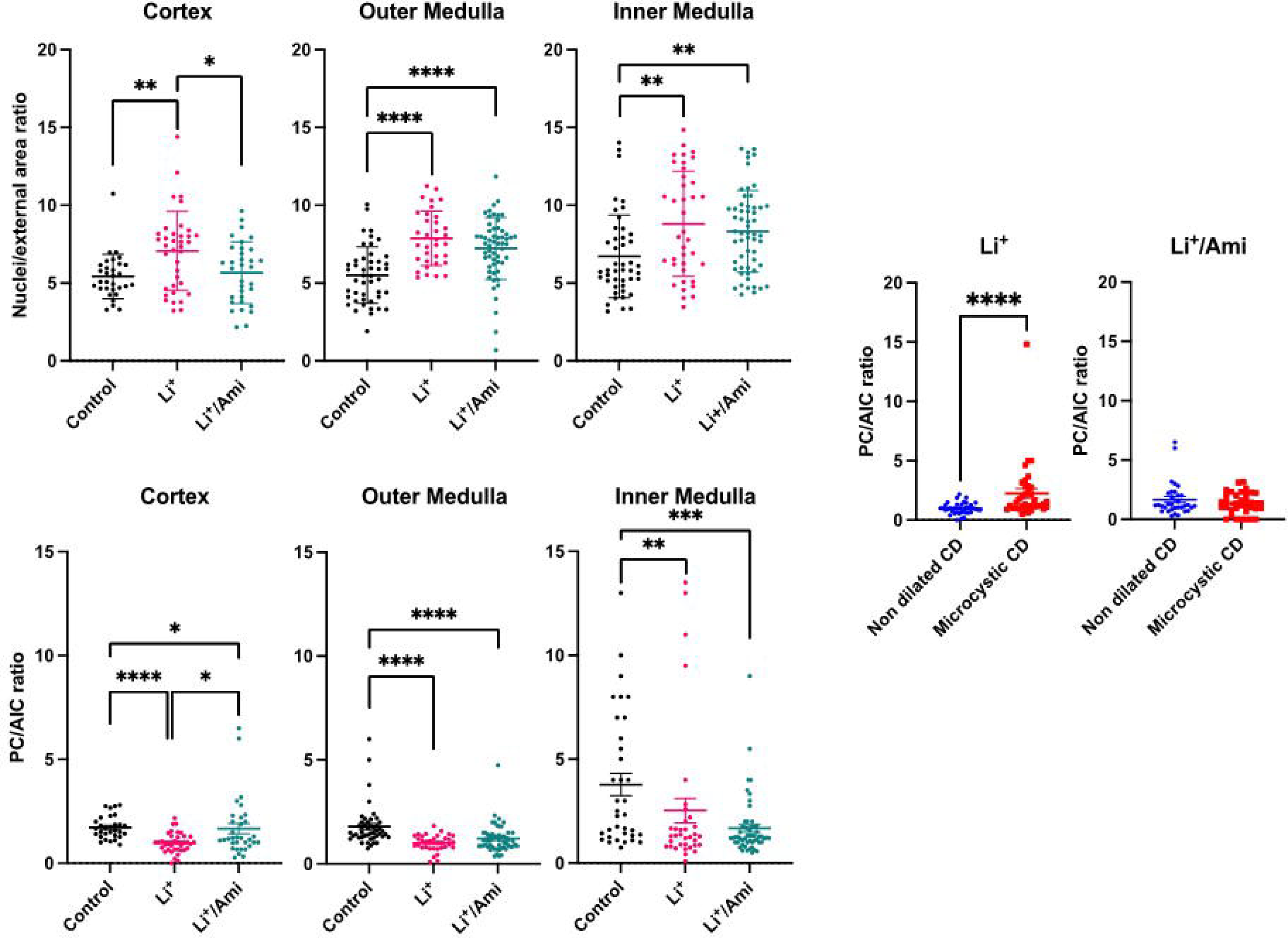
Masson trichrome staining of kidneys and quantification of tubular areas and minimal tubular diameters. Representative images of trichrome staining (left), automated tubular areas, and minimal diameters in each group (right). Only significant differences are represented. **** p < 0.0001. Uosm, urine osmolality; Li^+^, lithium; Li^+^/Ami, lithium and amiloride treatment.

### 3.2. A differential cell composition of CDs was observed in dilated and non-dilated CDs

Dilated tubules stained for markers of PCs and AICs, demonstrating that the tubular segment affected by dilations were CDs (Fig. 3). Morphometric analysis of dilated CDs showed an increase in the number of nuclei per tubular area in lithium-treated rats compared to controls, suggesting epithelial hypertrophy. This was equally observed in the cortex, the outer medulla, and the inner medulla (Fig. 4). Hypertrophy was not associated with positive Ki67 staining of CD cells at 6 months (Fig. 3).

**Fig. 3.**
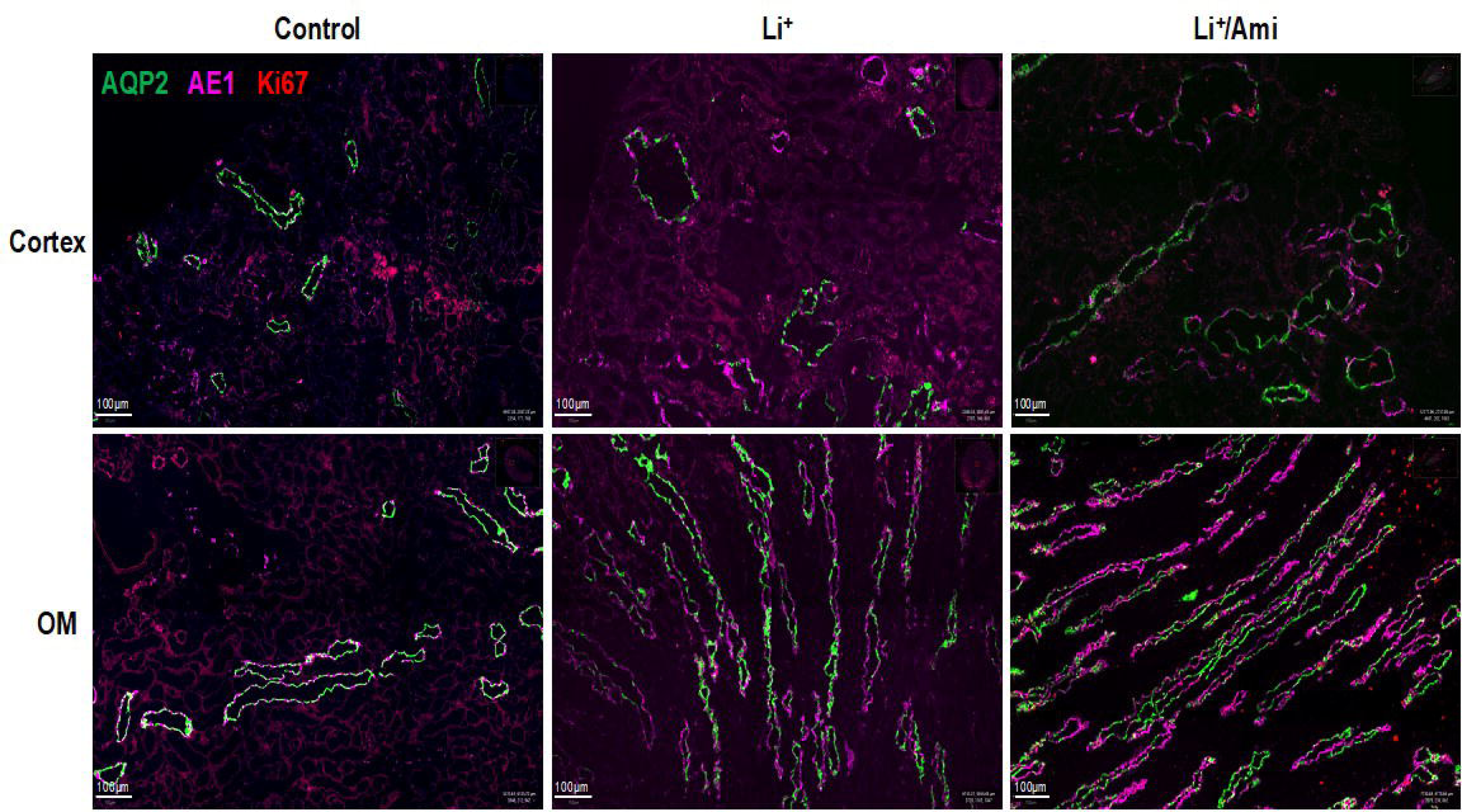
Immunofluorescent staining of collecting duct cells. Representative images of the cortex and the outer medulla (OM) of immunofluorescent staining of type-2 aquaporin (AQP2), type-1 anion exchanger (AE1), and Ki67 of kidney slices of control rats, rats treated with lithium (Li^+^), and rats treated with lithium and amiloride (Li^+^/Ami). No collecting duct cells stained positive for Ki67.

**Fig. 4.**
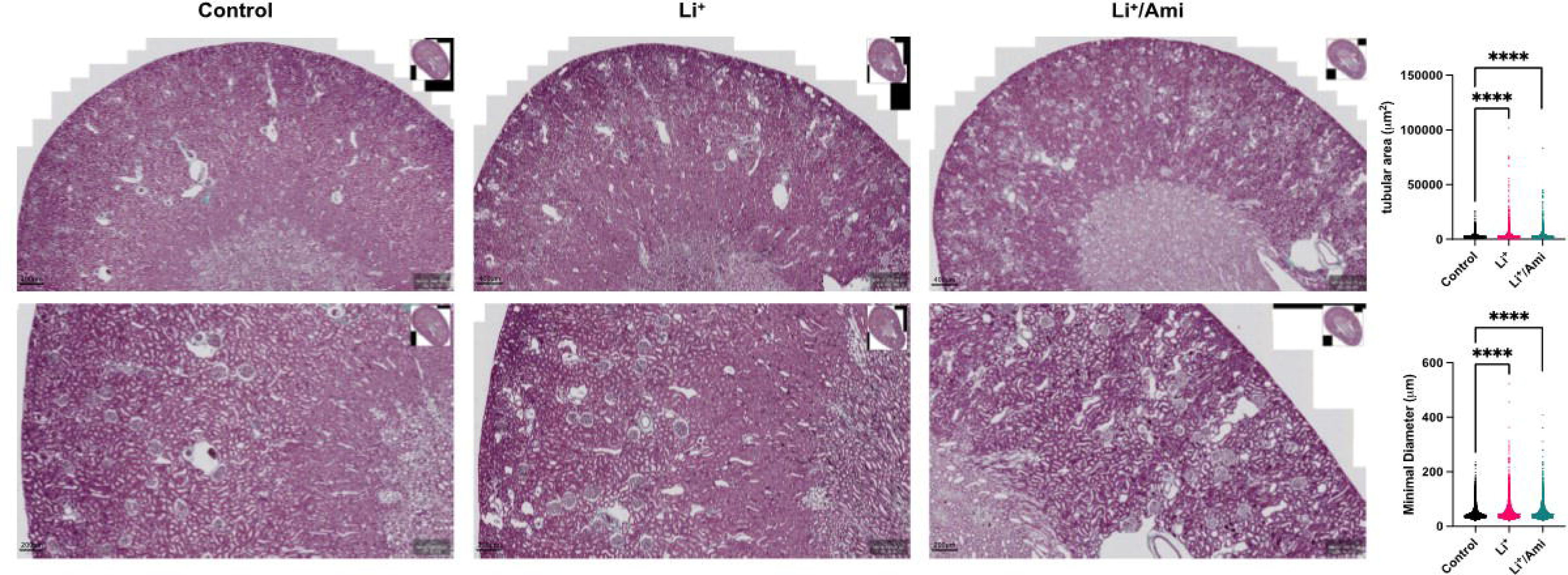
Morphometric analyses and cellular composition of collecting ducts. Upper panel, ratio between the numbers of DAPI-stained nuclei and external surface measurement of the collecting ducts in the cortex, the outer medulla and the inner medulla of control rats, rats treated with lithium (Li^+^), and rats treated with lithium and amiloride (Li^+^/Ami). Lower panel, ratio between principal cells (PC) and type-A intercalated cells (AIC) in the cortex, the outer medulla and the inner medulla of control rats, rats treated with lithium (Li^+^) and rats treated with lithium and amiloride (Li^+^/Ami). Right panel, ratio between principal cells (PC) and type-A intercalated cells (AIC) in the cortex of rats treated with lithium (Li^+^) and rats treated with lithium and amiloride (Li^+^/Ami) in non-dilated and dilated (microcystic) collecting ducts (CD). *P < 0.05, **P < 0.01, ***P < 0.001, ****P < 0.0001.

Analysis of CD composition revealed a decrease in PC-to-AIC ratio in the cortex and the outer and inner medulla of Li^+^ group rats (Fig. 4). However, cell composition of dilated CDs was different, lined with a majority of PCs, conversely to hypertrophied CDs (Fig. 4).

### 3.3. Despite no overt NDI, AQP2 expression decreased in lithium-treated rats despite weak correlation with cellular remodelling

AQP2 mRNA levels were analysed in the cortex, the outer stripe of the outer medulla (OSOM), the inner stripe of the outer medulla (ISOM), and the inner medulla, showing that mRNA levels were significantly lower in lithium-treated rats compared to controls in all these segments, despite the absence of significantly higher urine output, with decreased AQP2 expression gradient in the Li^+^ group (Fig. 5). There was no significant correlation between the cell and morphometric quantifications of CDs and the level of AQP2 expression except for a correlation between AQP2 expression and the PC-to-AIC ratio in the outer medulla (rho = 0.61, P = 0.03).

**Fig. 5.**
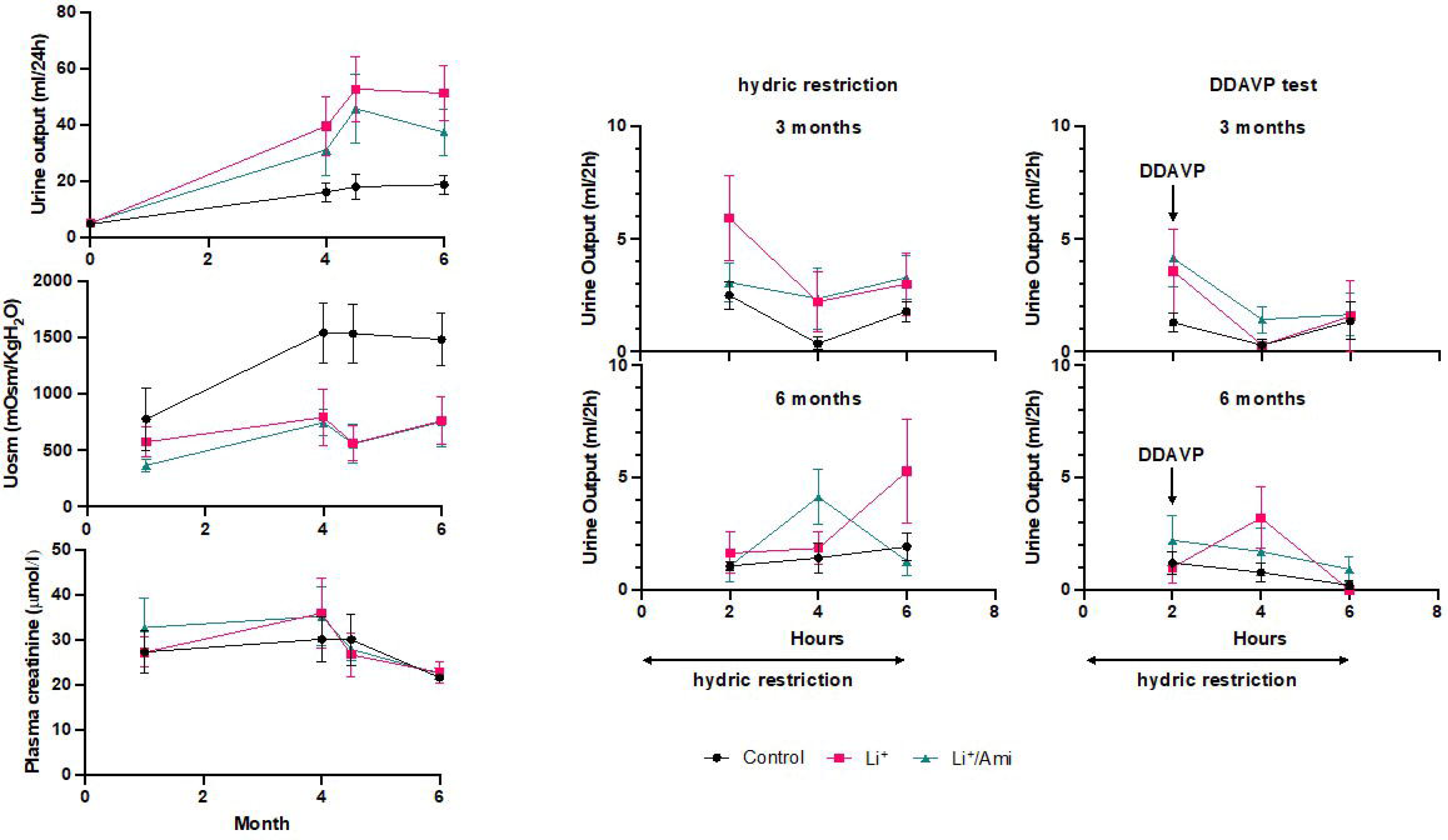
AQP2 expression in the different segments of the kidney. Relative AQP2 mRNA expression in the cortex, the outer stripe of the outer medulla (OSOM), the inner stripe of the inner medulla (ISOM), and the inner medulla (IM) in control rats, rats treated with lithium (Li^+^), and rats treated with lithium and amiloride (Li^+^/Ami). *P < 0.05, **P < 0.01, ***P < 0.001.

### 3.4. Amiloride treatment had no effect on microcystic dilations but decreased the PC-to-AIC ratio in the non-dilated portions of CDs

Plasma lithium concentration was 0.43 ± 0.16 mmol/L in the Li^+^/Ami group (P = 0.01, in comparison to the Li^+^ group). Urine output, urine osmolality, serum creatinine, and urine concentration ability after challenging tests did not differ between the Li^+^/Ami and Li^+^ groups (Fig. 1). Tubular area quantification did not also significantly differ (Fig. 2). Increased number of nuclei per external area was prevented exclusively in the renal cortex in the Li^+^/Ami group, with no significant differences in the outer or inner medulla (Fig. 3). This observation was associated with a less extended decrease in the PC-to-AIC ratio compared to the Li^+^ group (Fig. 3) but no difference in this ratio between dilated and non-dilated CDs (Fig. 4). Finally, AQP2 mRNA expression decrease was more marked, though not reaching statistical significance in all kidney segments in the Li^+^/Ami compared to the Li^+^ group (Fig. 5). Overall, amiloride did not prevent AQP2 transcription decrease after 6 months of lithium exposure.

## 4. Discussion

Our study showed in a rat model of lithium exposure that lower plasma lithium concentrations did not result in overt NDI but in morphological changes and cell remodelling consisting in microcystic dilations and hypertrophy of CDs.

Usual targeted plasma lithium range in bipolar-disease patients is 0.6-1.2 mmol/L. However, several clinical trials suggested clinical efficacy at lower lithium doses, with steady-state concentrations of 0.4 mmol/L (Nolen et al. 2019). Our previous study evaluating vasopressin resistance and NDI in lithium-treated patients showed that the main determinant of urine output was daily lithium dose, directly affecting plasma lithium concentrations, independently of the treatment duration (Tabibzadeh et al. 2022). Our experimental findings in a model with low plasma lithium concentrations are consistent with the clinical findings as, even though a significant decrease in AQP2 expression was observed, no significant change in urinary output, urine osmolality or urine concentration ability were evidenced. Such changes are observed in rats receiving therapeutic lithium concentrations, with clinical signs starting around 0.5 mmol/L (Nielsen et al. 2006). Immunofluorescent staining of AQP2 showed no apparent decrease in fluorescence intensity between the groups allowing identifying PCs with this staining. This suggests a resistance to the action of vasopressin to a certain degree, which might be obvious during hydric restriction challenge.

Despite the absence of polyuria or major AQP2 expression decrease, morphological changes were observed. These morphological changes involved CDs, as suggested by previous studies but with no direct evidence yet (Markowitz et al. 2000; Walker et al. 2005).Two features coexisted in all analysed kidneys, namely CD hypertrophy, characterized by an increase in the number of cells per tubular area, and microcystic dilations, defined by CD enlargement and cellular stretching. Cell composition differed with a majority of PCs in dilated CDs and a higher proportion of AICs in hypertrophic CDs. While Bissler et al. (2019) reported that cysts developed in a murine model of tuberous sclerosis complex originated from AICs, cysts in experimental models of autosomal polycystic kidney disease (ADPKD) consisted both of PCs and ICs, with a major role of PCs in cyst development (de Lemos Barbosa et al. 2016; Sullivan et al. 1998). Of note, primary cilia dysfunction, only present in PCs in CDs, represents an essential promoter of renal cysts in ADPKD and various cystic kidney diseases (generically referred to as ciliopathies) (McConnachie et al. 2021).

Cell remodelling has been previously reported in the course of lithium treatment and acute kidney injury (Park et al. 2018). In our study, no proliferation was seen after 6 months of lithium treatment, as assessed by Ki67 staining. Cells also presumably remained differentiated, with no double positive cells found using immunofluorescence analysis (AQP2^+^/AE1^+^). The existence, time course, and mechanisms of CD cell proliferation on lithium therapy are still a matter of debate. While Christensen et al. (2006) have found proliferation among PCs, De Groot et al. (2014) reported a G2 arrest of the cell cycle in these cells and a shift towards a higher proportion of AIC, in consistence with our findings. This shift might be due to trans-differentiation of PCs into AICs rather than proliferation, a mechanism likely mediated by Notch signalling inactivation (Mukherjee et al. 2019). This trans-differentiation is not associated with cellular proliferation in physiological conditions and acute kidney damage (Park et al. 2018). Early events remain to be evaluated in our model to determine a potential trans-differentiation with or without proliferation signals. Interestingly, the remodelling phenomena were barely correlated with the decrease in AQP2 expression, whereas such a correlation based on the decrease in PC proportion could be hypothesized. Our findings rather suggested that lithium-related effects on urine concentration and cell remodelling are uncoupled. Consistent, our previous reports evaluating lithium-treated bipolar disorder patients showed differential determinants of lithium-associated kidney texture changes using kidney magnetic resonance imaging radiomic analyses and NDI (Beunon et al. 2022; Tabibzadeh et al. 2022).

We did not evidence elevated serum creatinine concentrations or renal interstitial fibrosis suggestive of decrease in renal function after 6 months of lithium treatment, inconsistent with Walker et al. (2014), who reported interstitial fibrosis after the same duration of treatment but with a higher exposure to lithium. Whether cellular remodelling is deleterious in the long-term remains to be determined. However, our results, put in the perspective of previous literature, suggest that lowering kidney exposure to lithium might be nephroprotective only in a certain extent.

Finally, we studied the effect of amiloride treatment in this experimental under-exposed model. Urine concentration did not differ between the Li^+^ and Li^+^/Ami groups. AQP2 mRNA expression was even lower, although not significantly, in the Li^+^/Ami group. However, this should be interpreted with caution, as plasma lithium concentrations were higher, though still at infra-therapeutic levels, than in the Li^+^ group. As a monovalent cation, lithium enters tubular cells using the same transporters as sodium. As a specific ENaC blocker, amiloride has been proven to inhibit lithium entry in PCs *in vitro* and *in vivo* (Kortenoeven et al. 2009). Conversely, Bedford et al. demonstrated improved urine concentration and increased AQP2 expression when lithium-exposed rats were treated with amiloride (Bedford et al. 2008). Small clinical trials have shown an increase in urine osmolality and urinary AQP2 excretion after treating patients with amiloride (Battle et al. 1985; Bedford et al. 2008). In a long term (5 months) murine model of lithium-related nephrotoxicity, Kalita-De Croft et al. (2018) also showed that amiloride prevented renal fibrosis. The difference in lithium exposure compared to previous literature might explain the differences in our results. Moreover, in our study, increased lithium concentrations in the Li+/Ami group compared to the Li+ group were surprising, as amiloride is supposed to inhibit lithium reabsorption. These higher concentrations may therefore explain why we did not observe more preventing effects of amiloride. A mild effect of amiloride was observed on cellular remodelling, in particular in the renal cortex, with a lesser decrease in the PC-to-AIC ratio. This effect is in line with recent literature showing an important crosstalk between PCs and AICs, especially during acidosis (Guetin et al. 2013; Kleyman et al. 2013; Cheval et al. 2021; Tabibzadeh and Crambert 2022).

Our study has limitations. First, our design lacked an experimental group with plasma lithium concentrations in the therapeutic range, further useful to investigate the effects of amiloride on hypertrophy and microcysts development. Second, our study also lacked early time point assessments to evaluate early biochemical and morphological changes induced by exposure to lower plasma lithium concentrations. However, our data represent an interesting complement to previous studies on exposure to pharmacological lithium concentrations, with a model showing uncoupling of NDI and microcystic dilations, able to differentiate the involved mechanisms. Further studies are warranted to better evaluate CD remodelling in this setting.

In conclusion, our study demonstrated that under-exposure of kidneys to lithium results in the prevention of overt nephrogenic diabetes insipidus despite decrease in AQP2 expression, but did not prevent CD hypertrophy and microcysts, with only minor effects of amiloride on cell remodelling. Future studies are needed to clarify the exact signalling pathways leading to ICA hypertrophy and proliferation on one hand and microcysts generated from PCs on the other hand. Dose-effects studies should be helpful to better understand the onset of the different adverse kidney effects on lithium, even at low concentrations.

## Declarations

### Data Availability statement

The data that support the findings of this study are available from the corresponding author upon reasonable request. Some data may not be made available because of privacy or ethical restrictions.

### Institutional Review Board Statement

The animal study protocol was approved by the Ethics committee of Université Paris Cité and by the French Ministry of Research (approval # 26674-2020020714451509).

### Author contributions

**Nahid Tabibzadeh:** conceptualization; methodology; software; validation; formal analysis; investigation; resources; data curation; supervision; project administration; writing-original draft preparation; writing-review and editing. **Mathieu Klein:** methodology; validation; formal analysis; investigation; resources; data curation; writing-original draft preparation; writing-review and editing. **Mélanie Try:** methodology; software; validation; formal analysis; investigation; resources; data curation; writing-review and editing. **Joël Poupon:** validation; investigation; resources; writing-review and editing. **Pascal Houillier:** validation; investigation; resources; writing-review and editing. **Christophe Klein:** software; validation; investigation; resources; writing-review and editing. **Lydie Cheval:** validation; investigation; resources; writing-review and editing. **Gilles Crambert:** validation; investigation; resources; writing-review and editing. **Samia Lasaad:** validation; investigation; resources; writing-review and editing. **Lucie Chevillard:** conceptualization; methodology; validation; formal analysis; investigation; resources; supervision; project administration; writing-review and editing. **Bruno Megarbane:** conceptualization; methodology; validation; investigation; resources; supervision; project administration; funding acquisition; writing-review and editing.

